# Evidence for a reactionary account of retrieval state initiation

**DOI:** 10.1101/2025.01.28.635383

**Authors:** Subin Han, Nicole M. Long

**Affiliations:** Department of Psychology, University of Virginia

**Keywords:** memory, encoding, retrieval, EEG, attention, mvpa

## Abstract

Engagement of the retrieval state (or mode) is theorized to be a precursor to successful retrieval, but precisely when the retrieval state is engaged is unclear. Our aim in the present study is to determine the time course of retrieval state initiation. Based on growing evidence that the retrieval state reflects internal attention, we hypothesize that the retrieval state is reactionary rather than preparatory, whereby the retrieval state is engaged following – rather than preceding – a stimulus. We collected scalp electroencephalography data during a mnemonic state task in which we explicitly biased participants to engage in either retrieval or encoding states, and in which we manipulated the stimulus onset asynchrony (SOA) between the instruction and stimulus onsets. Our general expectation is that selective engagement of the retrieval state will only occur after stimulus presentation and thus be unaffected by the SOA. Our behavioral results show that regardless of the SOA, a bias to retrieval compared to encoding leads to worse memory for object stimuli. Using cross-study multivariate pattern analyses, we find robust engagement of the retrieval state approximately 500 ms following stimulus onset and no evidence for selective retrieval state engagement during the instruction interval. Together, these findings suggest that retrieval state engagement is reactionary and is consistent with the retrieval state reflecting internal attention engaged in the attempt to access stored information.

## 1 Introduction

Accessing stored episodic information underlies a host of behaviors and allows us to effectively engage with the world. For instance, as you plan dinner tonight, you may need retrieve what you had for dinner last night. However, how we initiate this access process remains a critical open question. According to one view, a retrieval state (or “mode;” Tulving, 1983) supports episodic remembering whereby one intentionally initiates a retrieval state in response to an instruction to retrieve. Initiation of the retrieval state enables you to treat the stimulus – e.g. “dinner” – as a retrieval cue in order to access stored episodic information. However, insofar as the retrieval state reflects internal attention (Chun, Golomb, & Turk-Browne, 2011; Long, Kuhl, & Chun, 2018; Long, 2023) and a target is required for internal attention (Chun et al., 2011), recruitment of the retrieval state should only occur after, not before, the stimulus appears. That is, rather than initiate a retrieval state, use “dinner” as a cue, and then attempt to access a stored representation, it is only after the target “dinner” is specified that you initiate the retrieval state, turning your mind’s eye inward to access what you had for dinner last night. Thus, whether we initiate retrieval prior to or following a stimulus remains an open question. The aim of this study was to determine the time course of retrieval initiation as the time course of retrieval is expected to impact downstream processing and behavioral outcomes.

The retrieval state may be preparatory, activated in response to an instruction to prepare to engage in episodic retrieval (Rugg & Wilding, 2000). Early work compared electrophysiological signals when participants were instructed to prepare to make a recognition (episodic) judgment vs. a semantic judgment (Morcom & Rugg, 2002) and found that approximately 600 ms after this instruction, a right frontal positivity emerged in response to the episodic vs. semantic instruction. Subsequent work has found a similar signal when contrasting instructions to prepare for episodic retrieval vs. a perceptual task in which no retrieval is required (Evans, Williams, & Wilding, 2015). The interpretation of these findings is that the retrieval state serves as a precursor to, but is separate from, episodic retrieval success. The individual first engages the retrieval state which enables them to treat the probe stimulus as an episodic retrieval cue allowing the search process to begin which may or may not lead to access of the stored representation.

Recent work, however, suggests that the retrieval state may instead be reactionary. Using multivariate pattern analysis of spectral signals across the scalp, we have found that we can distinguish between retrieval and encoding states around 1000 ms following *stimulus* onset (Long & Kuhl, 2019; Smith, Moore, & Long, 2022). As a preparatory retrieval state would be engaged prior to the stimulus, we would have anticipated robust memory state dissociations throughout the entirety of the stimulus interval. However, these late stimulus interval findings are in line with growing evidence from behavioral, neural and computational investigations (Logan, Cox, Annis, & Lindsey, 2021; Long, 2023; Logan et al., 2024) which suggest that the retrieval state may reflect internal attention. Whereas external attention is a focus on sensory, perceptual information, internal attention is the directing of the mind’s eye inward to stored representations and task goals (Chun et al., 2011). In this framework, a target is needed upon which to focus internal attention which is consistent with memory state dissociations following, rather than preceding, stimulus onset.

Whether retrieval state initiation is preparatory or reactionary is expected to have consequences for episodic remembering. Engaging the retrieval state both impacts how stimuli are processed (Long & Kuhl, 2021) and trades off with engaging the encoding state (Hasselmo, Bodelon, & Wyble, 2002; Long & Kuhl, 2019) such that retrieving instead of encoding can negatively impact subsequent memory (Long & Kuhl, 2019). The time course of retrieval state initiation will clarify how episodic remembering unfolds and how probe stimuli are processed to support successful remembering. Insights into the timing of retrieval initiation will additionally inform our understanding of the relationship between mnemonic brain states and attentional processing.

Our hypothesis is that the retrieval state is reactionary, and sustained over time. Alternatively, the retrieval state may be preparatory – that is, engaged following the instruction to retrieve and prior to stimulus onset. To test our hypothesis, we conducted a mnemonic state task in which participants studied two sets of object images while recording scalp electroencephalography (EEG). The second set of object images categorically overlapped with the first set, and during the second set, participants were explicitly instructed to either encode the second (present) object or retrieve the first (past) object. These instructions were intended to bias participants toward either an encoding or retrieval state. Our critical manipulation was the stimulus onset asynchrony (SOA) between the instruction and the stimulus onsets, whereby the shorter the SOA, the shorter the time between the instruction and the stimulus. We used cross-study multivariate pattern analyses (MVPA) to measure retrieval state evidence (i.e., evidence for retrieval state engagement) over time. To the extent that the retrieval state is reactionary and engaged following stimulus onset, we should find selective engagement of the retrieval state based on the instruction to encode or retrieve following stimulus onset for all SOA conditions.

## 2 Materials and Methods

### 2.1 Participants

Forty (28 female; age range = 18-54 years, mean age = 25.6 years) English speakers from the University of Virginia community participated. This sample size is based on our prior work in which we find an effect of mnemonic state instructions on recognition memory (Smith et al., 2022) as described in the preregistration report of this study (https://osf.io/xnwg9). All participants had normal or corrected-to-normal vision. Informed consent was obtained in accordance with the University of Virginia Institutional Review Board for Social and Behavioral Research, and participants were compensated for their participation. One participant was excluded from the final dataset due to technical issues resulting in poor signal quality throughout the majority of the session. Thus, data are reported for the remaining 39 participants.

### 2.2 Mnemonic state task experimental design

The task design follows our previous work (Smith et al., 2022) with one critical manipulation. Stimuli consisted of 864 object pictures, drawn from an image database with multiple exemplars per object category (Konkle, Brady, Alvarez, & Aude, 2010). From this database, we chose 216 unique object categories and four exemplars from each category. For each participant, one exemplar in a set of four served as a List 1 object, one as a List 2 object, and the two remaining exemplars served as lures for the recognition phase. Object condition assignment was randomly generated for each participant.

#### General overview

In each of twelve runs, participants viewed two lists, each containing 18 object images. For the first list (List 1), each object was new. For the second list (List 2), each object was again new but was categorically related to an object from the first list. For example, if List 1 contained an image of a bench, List 2 would contain an image of a different bench (Figure 1A). During List 1, participants were instructed to encode each new object. During List 2, however, each trial contained an instruction to either encode the current object (e.g., the new bench) or to retrieve the corresponding object from List 1 (the old bench). The critical manipulation was the stimulus onset asynchrony (SOA) between the instruction and the stimulus onset. SOA ranged from 500 ms to 2000 ms in 500 ms increments (Figure 1B) and varied across runs meaning that all List 2 trials within a run had the same duration. SOA conditions were randomly assigned across runs for each participant. Following twelve runs, participants completed a two-alternative forced-choice recognition test that separately assessed memory for all List 1 and List 2 objects.

**Figure 1.**
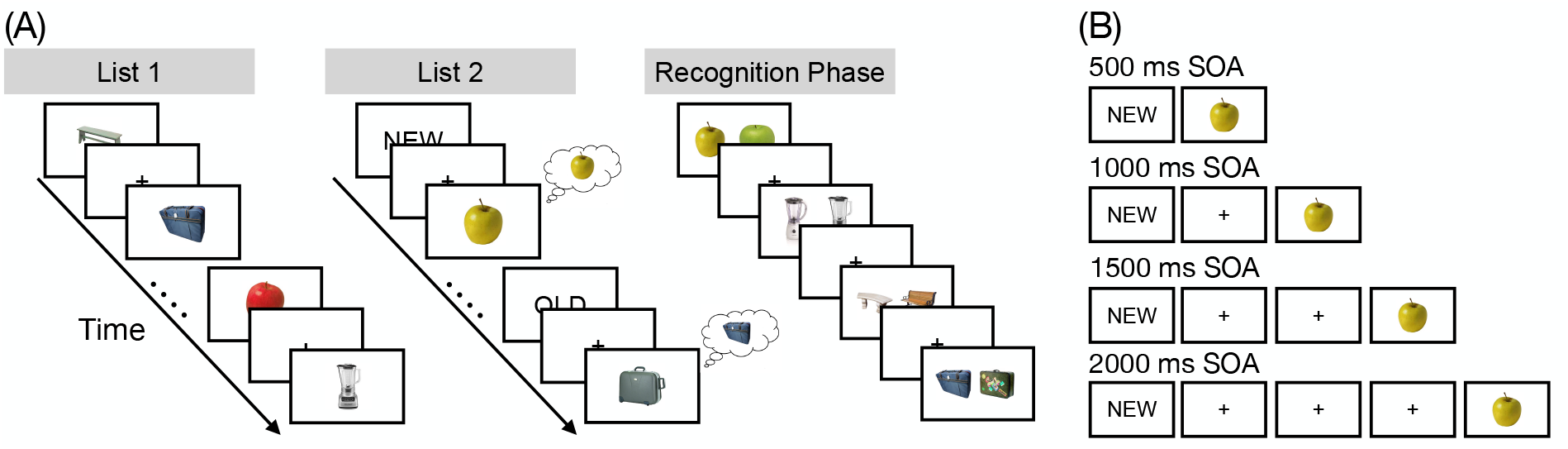
Task design. **(A)** During List 1, participants studied individual objects (e.g., apple, suitcase). During List 2, participants saw novel objects that were from the same categories as the objects shown in List 1 (e.g., a new apple, a new suitcase). Preceding each List 2 object was an OLD instruction cue or a NEW instruction cue. The OLD cue signaled that participants were to retrieve the corresponding object from List 1 (e.g., the old suitcase). The NEW cue signaled that participants were to encode the current object (e.g., the new apple). Each run of the experiment contained a List 1 and List 2; object categories (e.g., apple) were not repeated across runs. During List 2 trials, we manipulated stimulus onset asynchrony (SOA) between the instruction onset and the stimulus onset. SOA varied across runs meaning that all List 2 trials within a run had the same SOA. After twelve runs, participants completed a two-alternative forced-choice recognition test that tested memory for each List 1 and List 2 object. On each trial, a previously presented object, either from List 1 or List 2, was shown alongside a novel lure from the same category. Participants’ task was to choose the previously presented object. List 1 and List 2 objects were never presented together. **(B)** There were four SOA conditions. The shortest SOA was 500 ms and the longest SOA was 2000 ms. The “NEW” or “OLD” instruction cue was presented for 500 ms. Either immediately following the cue (SOA = 500 ms) or after a delay of 500, 1000, or 1500 ms during which a fixation cross was shown (SOAs *>* 500 ms), an object image was presented for 2000 ms.

#### List 1

On each trial, participants saw a single object presented for 2000 ms followed by a 1000 ms interstimulus interval. participants were instructed to study the presented object in anticipation of a later memory test.

#### List 2

On each trial, participants saw a cue word, either “OLD” or “NEW” for 500 ms. Either immediately following the cue (SOA = 500 ms) or after a fixation cross (SOAs *>* 500 ms), an object image was presented for 2000 ms (Figure 1B). All objects in List 2 were nonidentical exemplars drawn from the same category as the objects presented in the immediately preceding List 1. That is, if a participant saw a bench and an apple during List 1, a different bench and a different apple would be presented during List 2. On trials with a NEW instruction (encode trials), participants were to encode the presented object. On trials with an OLD instruction (retrieve trials), participants tried to retrieve the categorically related object from the preceding List 1. Importantly, this design prevented participants from completely ignoring List 2 objects following OLD instructions in that they could only identify the to-be-retrieved object category by processing the List 2 object.

Participants completed twelve runs with two lists in each run (List 1, List 2). Participants viewed 18 objects per list, yielding a total of 432 object stimuli from 216 unique object categories. Participants did not make a behavioral response during either List 1 or 2. Following the twelve runs, participants completed a two-alternative forced-choice recognition test.

#### Recognition phase

Following the twelve runs, participants completed the recognition phase. On each trial, participants saw two exemplars from the same object category (e.g., two benches; Figure 1A). One object had previously been encountered either during List 1 or List 2. The other object was a lure and had not been presented during the experiment. Because both test probes were from the same object category, participants could not rely on familiarity or gist-level information to make their response (Brainerd & Reyna, 2002). Trials were self-paced, and participants selected (via button press) the previously presented object. Trials were separated by a 1000 ms interstimulus interval. There were a total of 432 recognition trials (corresponding to the 432 total List 1 and List 2 objects presented in the experiment). List 1 and List 2 objects never appeared in the same trial together; thus, participants never had to choose between two previously presented objects. List 1 and List 2 objects were presented randomly throughout the test phase.

### 2.3 EEG data acquisition and preprocessing

All acquisition and preprocessing methods are based on our previous work (Smith et al., 2022); for clarity we use the same text as previously reported. EEG recordings were collected using a BrainVision system and an ActiCap equipped with 64 Ag/AgCl active electrodes positioned according to the extended 10-20 system. All electrodes were digitized at a sampling rate of 1000 Hz and were referenced to electrode FCz. Offline, electrodes were later converted to an average reference. Impedances of all electrodes were kept below 50 kΩ. Electrodes that demonstrated high impedance or poor contact with the scalp were excluded from the average reference; however, these electrodes were included in all subsequent analysis steps. Thus no electrodes and no data are excluded for any participant. Bad electrodes were determined by voltage thresholding (see below).

Custom python codes were used to process the EEG data. We applied a high pass filter at 0.1 Hz, followed by a notch filter at 60 Hz and harmonics of 60 Hz to each participant’s raw EEG data. We then performed three preprocessing steps (Nolan, Whelan, & Reilly, 2010) to identify electrodes with severe artifacts. First, we calculated the mean correlation between each electrode and all other electrodes as electrodes should be moderately correlated with other electrodes due to volume conduction. We z-scored these means across electrodes and rejected electrodes with z-scores less than -3. Second, we calculated the variance for each electrode, as electrodes with very high or low variance across a session are likely dominated by noise or have poor contact with the scalp. We then z-scored variance across electrodes and rejected electrodes with a |z| *>*= 3. Finally, we expect many electrical signals to be autocorrelated, but signals generated by the brain versus noise are likely to have different forms of autocorrelation. Therefore, we calculated the Hurst exponent, a measure of long-range autocorrelation, for each electrode and rejected electrodes with a |z| *>*= 3. Electrodes marked as bad by this procedure were excluded from the average re-reference. We then calculated the average voltage across all remaining electrodes at each time sample and re-referenced the data by subtracting the average voltage from the filtered EEG data. We used wavelet-enhanced independent component analysis (Castellanos & Makarov, 2006) to remove artifacts from eyeblinks and saccades.

### 2.4 EEG data analysis

We applied the Morlet wavelet transform (wave number 6) to the entire EEG time series across electrodes, for each of 46 logarithmically spaced frequencies (2-100 Hz; Smith et al., 2022). After log-transforming the power, we downsampled the data by taking a moving average across 100 ms time intervals from 4000 ms preceding to 4000 ms following object item presentation, and sliding the window every 25 ms, resulting in 317 time intervals (80 non-overlapping). Power values were then z-transformed by subtracting the mean and dividing by the standard deviation power. Mean and standard deviation power were calculated across all trials and across time points for each frequency.

### 2.5 Pattern classification analyses

Pattern classification analyses were performed using penalized (L2) logistic regression implemented via the sklearn module (0.24.2) in Python and custom Python code (Smith & Long, 2024). For all classification analyses, classifier features were composed of spectral power across 63 electrodes and 46 frequencies. Before pattern classification analyses were performed, an additional round of z-scoring was performed across features (electrodes and frequencies) to eliminate trial-level differences in spectral power (Kuhl & Chun, 2014; Long & Kuhl, 2018; Smith et al., 2022). Therefore, mean univariate activity was matched precisely across all conditions and trial types. We extracted “classifier evidence,” a continuous value reflecting the logit-transformed probability that the classifier assigns the correct mnemonic label (encode, retrieve) for each trial. Classifier evidence is used as a trial-specific, continuous measure of mnemonic state information, which is used to assess the degree of retrieval evidence present on List 2 encode and retrieve trials during both the instruction interval (i.e., the interval between the instruction and stimulus onset) and the stimulus interval.

### 2.6 Cross-study memory state classification

To measure retrieval state engagement for the List 2 instruction and stimulus intervals, we conducted three stages of classification using the same methods as in our prior work (Smith & Long, 2024). First, we conducted within participant leave-one-run-out cross-validated classification (penalty parameter = 1) on all participants who completed the mnemonic state task (N = 103; see Hong, Moore, Smith, & Long, 2023 for details). The classifier was trained to distinguish encoding versus retrieval states based on spectral power averaged across the 2000 ms stimulus interval during List 2 trials. For each participant, we generated true and null classification accuracy values. We permuted condition labels (encode and retrieve) for 1000 iterations to generate a null distribution for each participant. Any participant whose true classification accuracy fell above the 90th percentile of their respective null distribution was selected for further analysis (N = 37; 28 female; age range = 18-42, mean age = 21.24 years). This thresholding reduces the degree to which noise contributes to the training data by excluding participants who do not have robust spectral dissociations between encode and retrieve trials. Second, we conducted leave-one-participant-out cross-validated classification (penalty parameter = 0.0001) on the selected participants to validate the mnemonic state classifier and obtained classification accuracy of 60% which is significantly above chance (*t* _36_ = 6.000, *p <* 0.0001), indicating that the cross-participant mnemonic state classifier is able to distinguish encoding and retrieval states. Finally, we applied the cross-participant mnemonic state classifier to the current data across both the stimulus and instruction intervals during List 2 trials, specifically in 100 ms intervals from 2000 ms preceding to 2000 ms following stimulus onset. We extracted classifier evidence, the logit-transformed probability that the classifier assigned a given List 2 trial a label of encoding or retrieval. This approach provides a trial-level estimate of memory state evidence across the List 2 instruction and stimulus intervals.

### 2.7 Statistical Analyses

We used repeated measures ANOVAs (rmANOVAs) and *t* -tests to assess the effect of instruction (encode, retrieve) and SOA on behavioral memory performance. Also, we used rmANOVAs and *t* -tests to assess the effect of instruction, SOA, and time interval on memory state evidence during both the stimulus and instruction interval. We applied multiple comparisons corrections for post-hoc tests (using False Discovery Rate correction).

## 3 Results

### 3.1 Memory accuracy is modulated by list and instruction, but not SOA

We first sought to replicate our prior finding that participants can shift between encoding and retrieval in a goal-directed manner (Long & Kuhl, 2019; Smith et al., 2022). We expected to find higher accuracy for list 2 items studied with the encode relative to the retrieve instruction given the assumption that encoding and retrieval trade off. We assessed recognition accuracy specifically for the 2000 ms SOA condition (Figure 2A) as this matches our prior work. Following our pre-registration, we conducted a 2 *×* 2 rmANOVA with factors of list (1, 2) and instruction (encode, retrieve) and recognition accuracy as the dependent variable. We found a significant main effect of list (*F* _1,38_ = 16.07, *p* = 0.0003, *η*_*p*_^2^ = 0.2972) driven by greater recognition accuracy for list 1 items (*M* = 0.7963, *SD* = 0.1097) compared to list 2 items (*M* = 0.7407, *SD* = 0.1103). We did not find a significant main effect of instruction (*F* _1,38_ = 1.104, *p* = 0.3001, *η*_*p*_^2^ = 0.0282). We did not find a significant interaction between list and instruction (*F* _1,38_ = 2.985, *p* = 0.0922, *η*_*p*_^2^ = 0.0728). Although we failed to find a significant list by instruction interaction as in our prior work, we expect that this is due to the reduced number of trials in the present analysis. With the SOA manipulation, the number of trials per condition (list crossed with instruction) is 27 (three runs, 18 trials per list per run, half encode/half retrieve) which is substantially lower than the trial count in our prior work (72). The list 2 dissociation, although not significant (*t* _38_ = 1.980, *p* = 0.055, *d* = 0.2908), follows our previous observation and current prediction whereby memory accuracy is numerically greater for encode (*M* = 0.7588, *SD* = 0.1219) relative to retrieve trials (*M* = 0.7227, *SD* = 0.1263).

**Figure 2.**
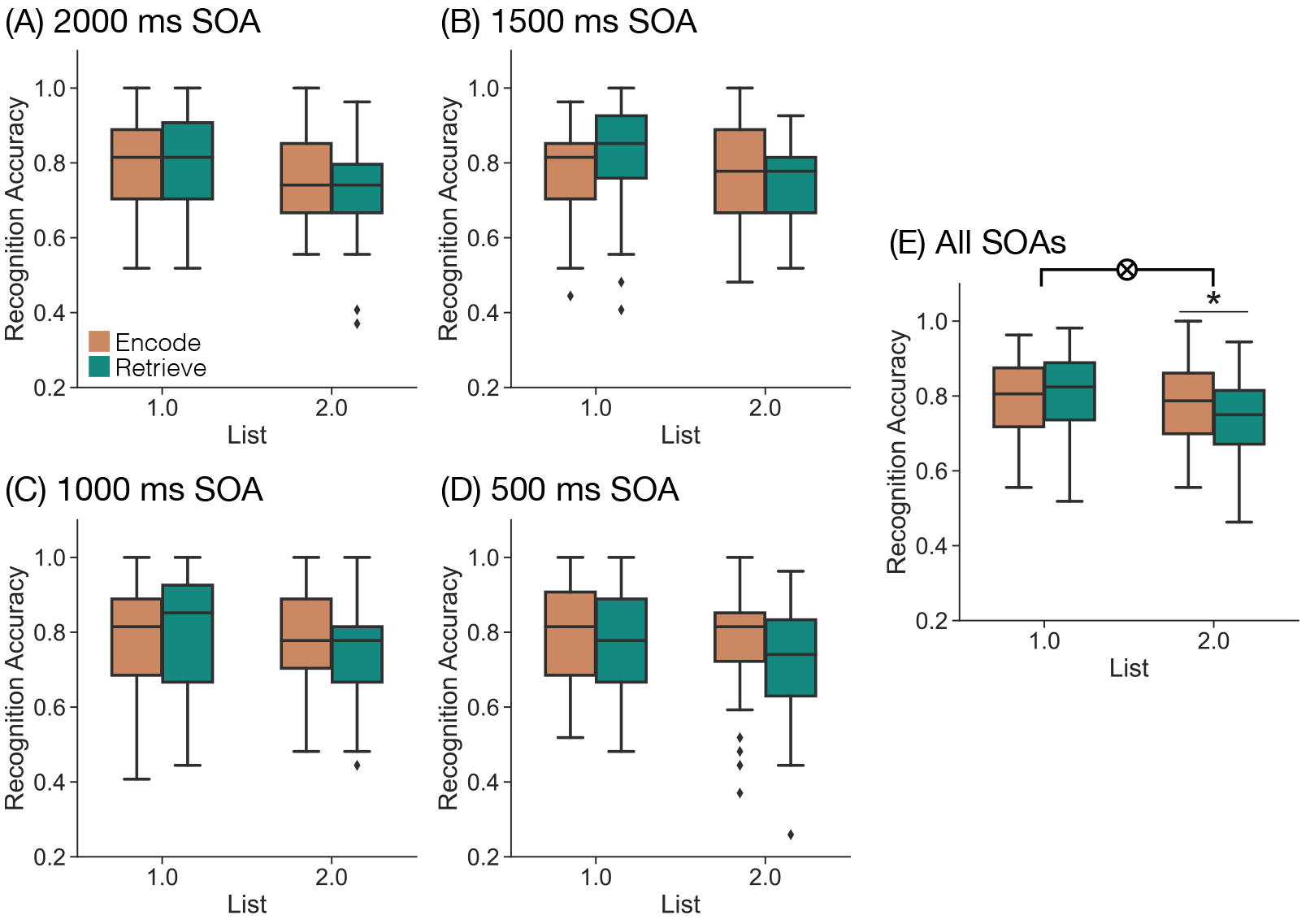
Influence of mnemonic instructions on memory behavior. We assessed recognition accuracy as a function of list (1, 2) and instruction (orange, encode; teal, retrieve) separately for each SOA condition (A-D) and for all trials averaged across SOA conditions (E). Box-and-whisker plots show the interquartile range, with median indicated by the horizontal line and whiskers denoting maximum and minimum (mean +/-2.5 x standard deviation). Diamonds indicate outliers. **(A-D)** We did not find any effect of SOA on recognition accuracy. **(E)** We found a significant interaction between list and instruction (p = 0.0011) driven by greater accuracy for list 2 encode compared to list 2 retrieve trials. *p*<*0.05, FDR corrected.

Our hypothesis is that the retrieval state is reactionary, or engaged following stimulus onset. Thus, SOA should not modulate the list by instruction interaction. That is, participants should engage in encoding on trials with the encode instruction and in retrieval on trials with the retrieve instruction after the stimulus onset regardless of the length of the instruction-stimulus interval. However, to the extent that the retrieval state is preparatory, with a longer SOA, participants will have more time to selectively engage in each memory state prior to stimulus onset. Thus, with longer SOAs, participants should have greater engagement in the respective memory states which will result in a larger interaction between list and instruction on memory accuracy as SOA increases. To test these two alternatives and following our pre-registration, we conducted a 2 *×* 2 *×* 4 rmANOVA with factors of list (1, 2), instruction (encode, retrieve), and SOA (500 ms, 1000 ms, 1500 ms, 2000 ms) and recognition accuracy as the dependent variable (Figure 2). We found a significant main effect of list (*F* _1,38_ = 15.31, *p* = 0.0004, *η*_*p*_^2^ = 0.2872) driven by greater recognition accuracy for list 1 items (*M* = 0.7926, *SD* = 0.1145) compared to list 2 items (*M* = 0.7518, *SD* = 0.1049). We found a significant main effect of instruction (*F* _1,38_ = 5.113, *p* = 0.0296, *η*_*p*_^2^ = 0.1186) driven by greater recognition accuracy for encode (*M* = 0.7787, *SD* = 0.1046) compared to retrieve trials (*M* = 0.7657, *SD* = 0.1081). We did not find a significant main effect of SOA (*F* _3,114_ = 0.6709, *p* = 0.5716, *η*_*p*_^2^ = 0.0174). As we expected, we found a significant interaction between list and instruction (*F* _1,38_ = 12.52, *p* = 0.0011, *η*_*p*_^2^ = 0.2478). This interaction was driven by significantly greater recognition accuracy for list 2 encode (*M* = 0.7707, *SD* = 0.11) compared to list 2 retrieve trials (*M* = 0.7329, *SD* = 0.1074; *t* _38_ = 4.109, *p* = 0.0002, *d* = 0.3472, FDR corrected), and numerically greater recognition accuracy for list 1 retrieve (*M* = 0.7984, *SD* = 0.1232) compared to list 1 encode trials (*M* = 0.7868, *SD* = 0.1121; *t* _38_ = -1.3032, *p* = 0.2003, *d* = -0.0988). We did not find a significant interaction between either list and SOA (*F* _3,114_ = 0.6241, *p* = 0.6009, *η*_*p*_^2^ = 0.0162) or instruction and SOA (*F* _3,114_ = 0.7399, *p* = 0.5304, *η*_*p*_^2^ = 0.0191). We did not find a significant three-way interaction between list, instruction, and SOA (*F* _3,114_ = 0.3737, *p* = 0.7721, *η*_*p*_^2^ = 0.0097). Bayes factor analysis revealed that a model without the three-way interaction term (H_0_) is preferred to a model with the three-way interaction term (H_1_; H_10_ = 0.0146 very strong evidence for H_0_).

Together, our behavioral results support the reactionary account. We find robust evidence that participants can selectively engage in memory encoding and memory retrieval, replicating our past work. We do not find evidence that the amount of preparation time or SOA between instruction and stimulus influences the impact of list and instruction on memory accuracy.

### 3.2 Retrieval state evidence is modulated by instruction after stimulus onset, but not before

Our central goal was to test the hypothesis that the retrieval state is reactionary, and sustained over time. To test our hypothesis, we applied a cross-study mnemonic state classifier to trial-level spectral data across the whole brain, and leveraged the high temporal resolution of EEG and examined retrieval state evidence for 100 ms time intervals during the instruction and stimulus intervals. Specifically, we extracted retrieval state evidence from 500 ms to 2000 ms preceding (based on the SOA condition) to 2000 ms following the stimulus onset. To the extent that the retrieval state is reactionary and engaged after stimulus onset, we should find no dissociation in retrieval state evidence on the basis of instruction during the instruction interval and only find a dissociation during the stimulus interval. However, to the extent that the retrieval state is preparatory and engaged in response to the instruction cue, we should find dissociation in retrieval state evidence on the basis of instruction during the instruction interval. There should be greater dissociation with longer SOAs, and the dissociation should linger into the stimulus interval.

We first sought to replicate our prior finding that there is greater retrieval evidence on retrieve compared to encode trials later in the stimulus interval (Smith et al., 2022). For this analysis, we focused exclusively on the 2000 ms SOA condition given that we used a fixed 2000 ms SOA in our prior work. Following our pre-registration, we conducted a 2 *×* 20 rmANOVA with factors of instruction (encode, retrieve) and time interval (twenty 100 ms time intervals across the 2000 ms stimulus interval) and retrieval state evidence as the dependent variable (Figure 3A). We found a significant main effect of instruction (*F* _1,38_ = 10.56, *p* = 0.0024, *η*_*p*_^2^ = 0.2174) driven by greater retrieval evidence for retrieve (*M* = 0.0139, *SD* = 0.0621) compared to encode trials (*M* = -0.0286, *SD* = 0.0555). We found a significant main effect of time interval (*F* _19,722_ = 28.37, *p <* 0.0001, *η*_*p*_^2^ = 0.4274). We found a significant interaction between instruction and time interval (*F* _19,722_ = 2.392, *p* = 0.0008, *η*_*p*_^2^ = 0.0592) driven by greater retrieval evidence for retrieve compared to encode trials later in the stimulus interval (Figure 3A). Thus, we successfully replicated our prior finding that retrieval state evidence differs as a function of instruction during the stimulus interval.

**Figure 3.**
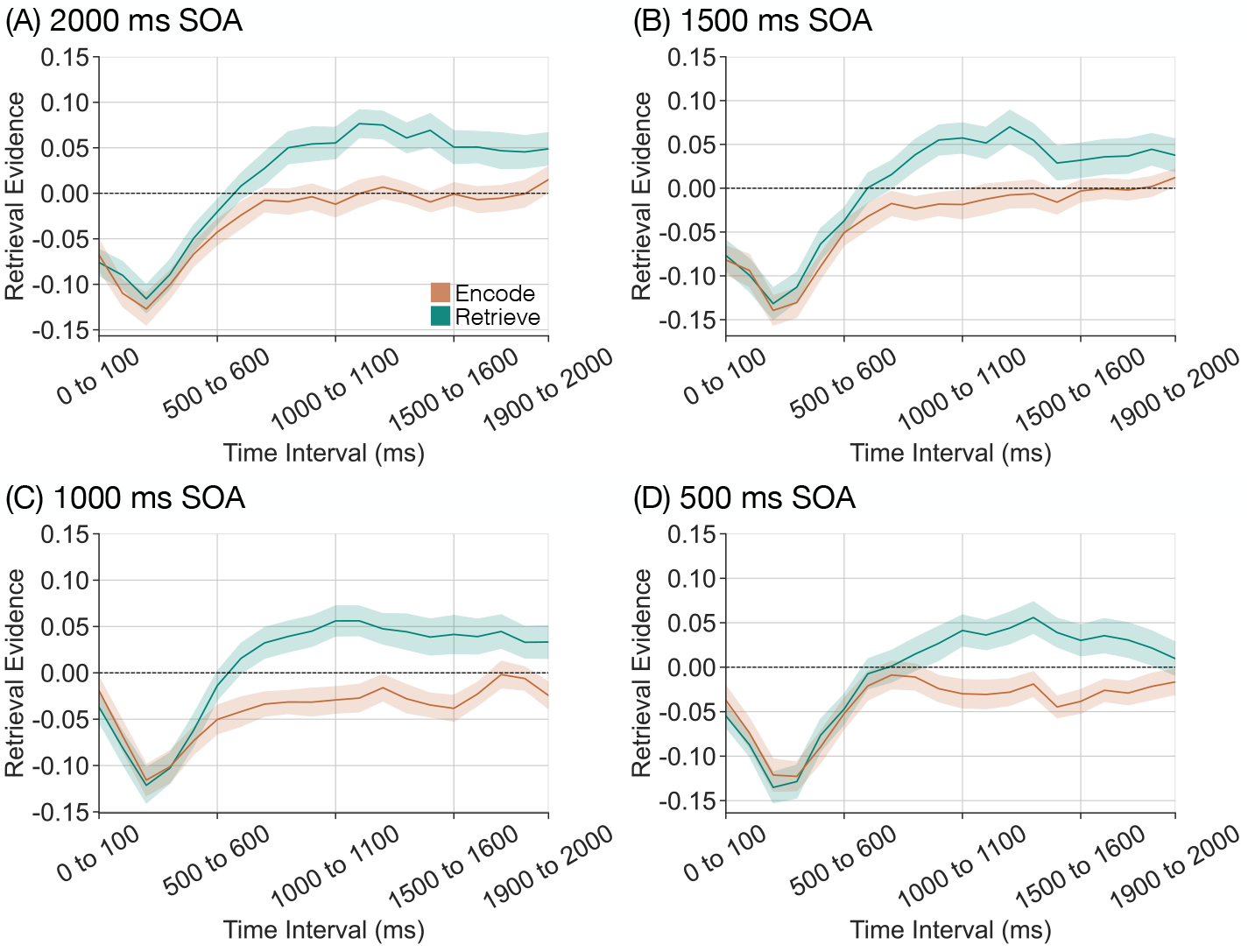
Influence of mnemonic instructions on stimulus-interval retrieval state evidence. We applied a cross-study classifier to the stimulus interval (twenty 100 ms time intervals) to measure retrieval state evidence as a function of instruction (encode, orange; retrieve, teal) separately for each stimulus onset asynchrony (SOA) condition. **(A-D)** Across all SOA conditions, we find a dissociation in retrieval state evidence on the basis of instruction around 500 ms after stimulus onset. Error bars reflect standard error of the mean.

Next, we evaluated the effect of SOA on retrieval state evidence during the stimulus interval. To the extent that the retrieval state is reactionary, we should find no effect of SOA on retrieval state evidence. However, to the extent that the retrieval state is preparatory, there should be greater instruction-interval dissociation in retrieval state evidence as a function of instruction as SOA increases. Thus, we should find an effect of SOA during the stimulus interval whereby there is a greater dissociation in retrieval state evidence for shorter SOA conditions compared to longer SOA conditions. Following our pre-registration, we conducted a 2 *×* 4 *×* 20 rmANOVA with factors of instruction (encode, retrieve), SOA (500 ms, 1000 ms, 1500 ms, 2000 ms), and time interval (twenty 100 ms time intervals) and retrieval state evidence as the dependent variable (Figure 3). We found a significant main effect of instruction (*F* _1,38_ = 15.23, *p* = 0.0004, *η*_*p*_^2^ = 0.2861) driven by greater retrieval evidence for retrieve (*M* = 0.0039, *SD* = 0.0537) compared to encode trials (*M* = -0.0368, *SD* = 0.04). We found a significant main effect of time interval (*F* _19,722_ = 45.30, *p <* 0.0001, *η*_*p*_^2^ = 0.5438). We did not find a significant main effect of SOA (*F* _3,114_ = 1.217, *p* = 0.3068, *η*_*p*_^2^= 0.031). We found a significant interaction between instruction and time interval (*F* _19,722_ = 9.9577, *p <* 0.0001, *η*_*p*_^2^ = 0.2076), and a significant interaction between SOA and time interval (*F* _57,2166_ = 1.602, *p* = 0.0031, *η*_*p*_^2^ = 0.0405). We did not find a significant interaction between instruction and SOA (*F* _3,114_ = 0.2892, *p* = 0.8331, *η*_*p*_^2^ = 0.0076). We did not find a significant three-way interaction between instruction, SOA, and time interval (*F* _57,2166_ = 0.6754, *p* = 0.9702, *η*_*p*_^2^ = 0.0175). Bayes factor analysis revealed that a model without the three-way interaction term (H_0_) is preferred to a model with the three-way interaction term (H_1_; H_10_ = 1.982*×*10^-8 extreme evidence for H_0_). In short, we found no effect of SOA on retrieval state evidence during the stimulus interval which supports the reactionary account whereby memory states are engaged following stimulus onset.

To investigate the extent to which the retrieval state is recruited during the instruction interval, we evaluated the effect of instruction on instruction-interval retrieval state evidence separately for each SOA condition (Figure 4). To the extent that the retrieval state is preparatory, we should find greater retrieval state evidence on retrieve compared to encode trials during the instruction interval. Following our preregistration, we conducted four separate rmANOVAs. Each ANOVA had factors of instruction (encode, retrieve) and time interval (from five to twenty 100 ms time intervals depending on the SOA condition) and retrieval state evidence as the dependent variable. We report the results of this ANOVA in Table 1 and highlight the key findings here.

**Table 1.** ANOVA results testing the effect of instruction and time interval on instruction-interval retrieval state evidence separately for each SOA condition.

| | Main effect of inst. | Main effect of time | Interaction of inst. × time |
| --- | --- | --- | --- |
| 500 ms |  |  |  |
| df | (1,38) | (4,152) | (4,152) |
| $F$ | 0.5128 | 13.477 | 2.033 |
| $p$ | 0.4783 | **<0.0001** | 0.0925 |
| $\eta_p^2$ | 0.0133 | 0.2618 | 0.0508 |
| 1000 ms |  |  |  |
| df | (1,38) | (9,342) | (9,342) |
| $F$ | 4.924 | 6.943 | 0.7532 |
| $p$ | **0.0325** | **<0.0001** | 0.6599 |
| $\eta_p^2$ | 0.1147 | 0.1545 | 0.0194 |
| 1500 ms |  |  |  |
| df | (1,38) | (14,532) | (14,532) |
| $F$ | 0.0794 | 8.050 | 0.5036 |
| $p$ | 0.7797 | **<0.0001** | 0.9315 |
| $\eta_p^2$ | 0.0021 | 0.1748 | 0.0131 |
| 2000 ms |  |  |  |
| df | (1,38) | (19,722) | (19,722) |
| $F$ | 0.0473 | 3.085 | 0.799 |
| $p$ | 0.8291 | **<0.0001** | 0.7098 |
| $\eta_p^2$ | 0.0012 | 0.0751 | 0.0206 |

Bold values indicate $p < 0.05$ .

**Figure 4.**
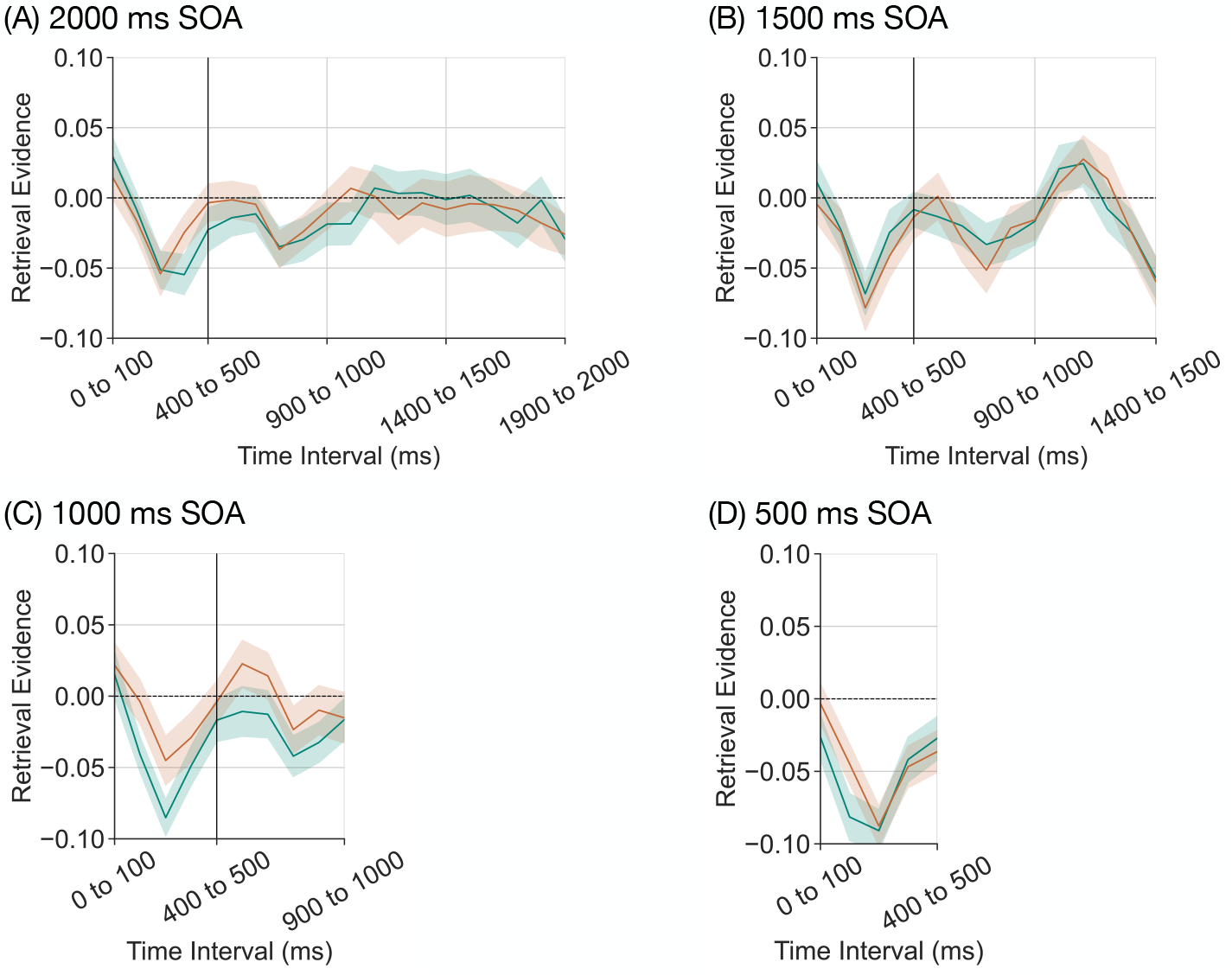
Influence of mnemonic instructions on instruction-interval retrieval state evidence. We applied a cross-study classifier to the instruction interval (five to twenty 100 ms time intervals depending on the SOA condition) to measure retrieval state evidence as a function of instruction (encode, orange; retrieve, teal) separately for each stimulus onset asynchrony (SOA) condition **(A-D)**. The vertical line at the 400 to 500 ms time interval indicates the offset of the instruction cue. Except for the 1000 ms SOA condition, we find no dissociation in retrieval evidence on the basis of instruction. **(C)** In the 1000 ms SOA condition we find greater retrieval evidence for encode compared to retrieve trials (*p* = 0.0325). Error bars reflect standard error of the mean.

Except for the 1000 ms SOA condition, we did not find a significant main effect of instruction, meaning that retrieval evidence did not significantly differ across retrieve vs. encode instructions. Bayes factor analysis revealed that a model without instruction (H_0_) is preferred to a model with instruction for the three SOA conditions (500 ms SOA: H_1_; H_10_ = 0.0263 very strong evidence for H_0_; 1500 ms SOA: H_1_; H_10_ = 2.730*×*10^-5 extreme evidence for H_0_; 2000 ms SOA: H_1_; H_10_ = 1.204*×*10^-5 extreme evidence for H_0_). Thus, we do not find evidence that participants selectively engage the retrieval state during the instruction interval. The significant effect that we found for the 1000 ms SOA condition was in the opposite direction as we predicted – retrieval state evidence was greater for encode (*M* = -0.0072, *SD* = 0.077) relative to retrieve trials (*M* = -0.0291, *SD* = 0.0604). Thus we failed to find evidence to support the interpretation that the retrieval state is preparatory.

Finally, we directly compared instruction-interval retrieval state evidence across SOA conditions. Based on prior work demonstrating that participants can engage control processes in preparation for an upcoming event (Gruber & Otten, 2010), we hypothesized that mnemonic state evidence might be modulated by SOA. Our classifier is designed such that less evidence for the retrieval state (negative evidence values) is equivalent to greater evidence for the encoding state. Insofar as the encoding state is engaged in anticipation of the upcoming stimulus, we expected that encoding state evidence would increase – or, conversely, that retrieval state evidence would decrease – as the SOA increased. Following our preregistration, we averaged retrieval state evidence across the instruction interval to account for the variable instruction interval length and conducted a 2 *×* 4 rmANOVA with factors of instruction (encode, retrieve) and SOA (500 ms, 1000 ms, 1500 ms, 2000 ms) and retrieval state evidence as the dependent variable.

We found a significant main effect of SOA (*F* _3,114_ = 3.957, *p* = 0.0100, *η*_*p*_^2^ = 0.0943). However, counter to our expectations, we found greater retrieval evidence for longer SOA compared to shorter SOA conditions (Figure 5). The main effect of instruction and the interaction between instruction and SOA were not significant (main effect: *F* _1,38_ = 1.730, *p* = 0.1963, *η*_*p*_^2^ = 0.0435; interaction: *F* _3,114_ = 1.072, *p* = 0.3639, *η*_*p*_^2^ = 0.0274). Together, these results suggest that participants may spend more time focused internally with longer SOAs.

**Figure 5.**
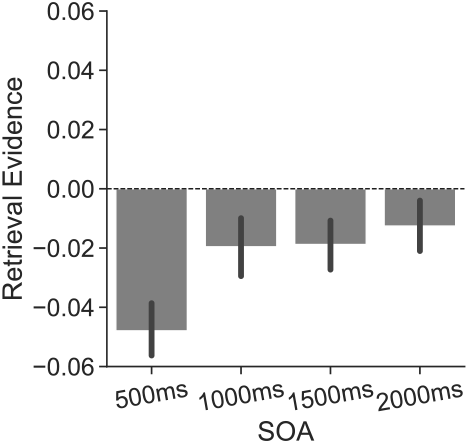
Influence of SOA on instruction-interval retrieval state evidence. We averaged retrieval state evidence across the instruction interval (five to twenty 100 ms time intervals depending on the SOA condition) and instruction (encode, retrieve) for each SOA condition. We find greater retrieval state evidence for longer SOA conditions. Error bars reflect standard error of the mean.

Together, our neural results support the reactionary account. We find robust evidence that participants selectively engage in memory encoding and memory retrieval after the stimulus onset. We do not find evidence that instruction influences retrieval state engagement during the instruction interval or that SOA affects the impact of instruction on retrieval state engagement during the stimulus interval. These results show that participants engage memory states following the stimulus onset, rather than in response to the instruction cue.

## Discussion

The goal of this study was to investigate the time course of retrieval initiation. Using cross-study decoding of scalp EEG data (Long, 2023), we measured retrieval state engagement during a task in which participants were explicitly instructed to either encode the present event or retrieve a past, overlapping event. We find behavioral evidence that participants’ ability to follow these instructions is unaffected by the stimulus onset asynchrony (SOA) between the instruction and stimulus. We find robust neural evidence that memory brain states are selectively engaged after stimulus onset, and that this selectivity is not modulated by SOA. Together, these findings suggest that the retrieval state may constitute internal attention which requires a target, proposing a link between memory brain states and attentional processes.

We find that the tendency to follow instructions to selectively encode vs. retrieve present vs. past stimuli is unaffected by the latency between the instruction and stimulus. Consistent with our prior work (Long & Kuhl, 2019; Smith et al., 2022), we find higher memory performance for currently presented items that are presented with an “encode” instruction relative to items presented with a “retrieve” instruction. These findings are in line with the idea that encoding and retrieval trade off (Hasselmo et al., 2002; Hasselmo, 2005) such that engaging in retrieval precludes engaging in encoding (Long & Kuhl, 2019). To the extent that participants engage mnemonic states in response to the instruction cue (Rugg & Wilding, 2000; Morcom & Rugg, 2002; Evans et al., 2015), we would anticipate a larger dissociation in memory performance with longer SOAs as a longer instruction-stimulus interval should provide additional time to recruit the goal-relevant state. However, we failed to find any effect of SOA and even for the shortest SOA, we find a dissociation in memory performance on the basis of instructions. Although we cannot draw strong conclusions from behavioral data alone, the robust behavioral effects across SOA suggests that mnemonic states are not selectively recruited during the instruction interval.

The retrieval state is consistently, selectively engaged during the stimulus interval regardless of SOA. A preparatory retrieval state should be engaged prior to stimulus onset (Morcom & Rugg, 2002; Evans et al., 2015) such that we should find retrieval state dissociations across all time points in the stimulus interval and to a greater degree for longer SOAs. That is, the magnitude of the difference in retrieval evidence between encode and retrieve trials should increase as SOA increases. However, we find robust retrieval state engagement in response to the retrieve instruction after stimulus onset for all SOAs. Such findings are inconsistent with preparatory mnemonic states, but do follow from our prior work in which we demonstrated retrieval state dissociations around 1000 ms after stimulus onset (Long & Kuhl, 2019; Smith et al., 2022). Furthermore, these findings are in line with the interpretation that the retrieval state reflects internal attention (Chun et al., 2011; Logan et al., 2021; Long, 2023; Smith & Long, 2024). According to this framework, episodic retrieval can be thought of as the process of turning the mind’s eye inward in the attempt to access a stored representation. A critical aspect of this framework is that a “target” is needed upon which attention can be focused – it is only possible to turn the mind’s eye inward once a target, in our study the visually presented stimulus, is provided. Thus, retrieval state dissociations should only be observed during the stimulus interval.

We find that retrieval state evidence during the instruction interval is modulated only by time and not by instruction. To the extent that the retrieval state is preparatory, we should have observed instruction-based dissociations whereby retrieval state evidence is greater for retrieve compared to encode trials and this effect should have been larger for longer SOAs. However, for all but one SOA, we failed to find an effect of instruction on retrieval evidence, and in the remaining SOA, the effect is in the opposite direction of what we would have predicted – namely, greater retrieval state evidence on *encode* rather than retrieve trials. Although inconsistent with the preparatory account, this finding is consistent with the interpretation that individuals engage internally directed attention in response to a target, rather than an instruction, and that the retrieval state reflects this internally directed attention. It is important to note that the mnemonic state classifier that we used here was, by design, trained on stimulus-interval data. We intentionally used this approach, rather than a separate instruction-interval classifier, as we expect there to exist a single, ubiquitous, retrieval state which if preparatory in nature, should be engaged during both the instruction and stimulus interval. Such an approach is in line with assumptions made in behavioral work demonstrating a stimulus-driven lingering influence of memory states (Duncan, Sadanand, & Davachi, 2012; Patil & Duncan, 2018). According to this work, hippocampally-mediated mnemonic states underlie the effect whereby responses on a prior trial influence memory accuracy on the current trial.

Our results appear to contradict past findings that demonstrate instruction-interval dissociations. Prior work on the retrieval state has demonstrated a positive voltage deflection over right prefrontal cortex approximately 600 ms following instruction onset (Morcom & Rugg, 2002). One potential avenue to reconcile these discrepancies is to consider the retrieval state in contrast to retrieval orientation. Whereas the retrieval state is characterized as a “tonic state that is maintained while episodic retrieval is required and remains constant across different episodic retrieval tasks” (Herron & Wilding, 2004), retrieval orientation should “vary according to the content of what episodic information is to be retrieved from memory” (Herron & Wilding, 2004). Extant studies have largely (though not exclusively, see Evans et al., 2015) compared episodic to semantic retrieval tasks on the premise that the retrieval state is specific to episodic memory. However, given the suggestion that the retrieval state may reflect internal attention, it should be recruited in the service of any internal attention demands, whether episodic or semantic (Buckner & DiNicola, 2019). Therefore, one intriguing possibility is that the episodic/semantic contrast is more akin to a superordinate retrieval orientation, rather than a retrieval state per se. This is an important direction for future work which we are actively pursuing. It is also important to consider that a preparatory retrieval state cannot fully account for evidence demonstrating automatic retrieval state engagement (Smith et al., 2022). For example, as you prepare tonight’s dinner of boeuf bourguignon, you may find yourself reminded of a visit to France in 2013, without ever having had the intention of retrieving said trip. A stimulus-driven retrieval state can account for the finding that temporally overlapping experiences induce the retrieval state independently from top-down demands (Smith et al., 2022). Additional consideration of the robustness of the right frontal positivity is also warranted, given that reports of this event-related potential are somewhat inconsistent. It has been selectively observed either only on “stay,” trials, when the instruction (e.g. “retrieve”) is the same as the prior trial (Morcom & Rugg, 2002) or only on “switch,” trials, when the instruction has changed from the prior trial (Evans et al., 2015). In theory, the retrieval state – insofar as it supports episodic remembering – should be recruited on both switch and stay trials. In the present study, we did not directly manipulate switch vs. stay trials and instead allowed instructions to randomly vary within a run such that participants will have encountered both types of trials. Even with this random variation, we find robust retrieval state engagement late in the stimulus interval.

Although we do not find an effect of instruction, we do find that SOA modulates instruction-interval retrieval state evidence. Specifically, we find that retrieval state evidence increases as a function of SOA. We have intentionally built a classifier which places encoding and retrieval on a continuum based on the hypothesis that encoding and retrieval tradeoff (Hasselmo et al., 2002; Hasselmo, 2005; Long & Kuhl, 2019; Smith et al., 2022). Thus, given the design of our classifier, less retrieval state evidence corresponds to greater encoding state evidence. This in turn means that we find greater encoding state evidence for shorter SOAs. Our interpretation is that encoding state evidence reflects *external* attention (Chun et al., 2011) – both to the instruction cue and to the stimulus. There is an initial increase in encoding evidence in the 500 ms following cue onset across all SOAs (Figure 4). For the shortest SOA, this is immediately followed by stimulus onset. In the longer SOAs, participants have more time (minimum 500 ms, maximium 1500 ms) during which only a fixation cross is presented. It may be that after the offset of the instruction, participants transition into a retrieval state – regardless of instruction – either to think about what the instruction means or maybe even to mind-wander or zone out (Mildner & Tamir, 2019; Esterman, Noonan, Rosenberg, & DeGutis, 2013).

In conclusion, our findings suggest that the retrieval state is reactionary and sustained over time. We find consistent, selective engagement of retrieval state evidence on the basis of instructions during the stimulus interval, but not during the instruction interval, across all SOA conditions. These findings suggest that the attempt to access stored representations occurs after stimulus onset and is consistent with prior findings (Logan et al., 2021; Long, 2023; Logan et al., 2024) which suggest that the retrieval state may constitute internal attention. Together, these findings advance our understanding of the relationship between attention and memory, and how retrieval may be initiated.

## Data and Code Availability

The datasets generated in the current study, along with all experimental codes used for data collection and data analysis will be made available on the Open Science Foundation (OSF) via the Long Term Memory Lab website (http://longtermmemorylab.com/publications/) upon publication.

## Author Contributions

Subin Han: Data curation; Formal analysis; Software; Visualization; Writing–Original draft; Writing– Review & editing. Nicole M. Long: Conceptualization; Formal analysis; Funding acquisition; Project administration; Software; Supervision; Visualization; Writing–Original draft; Writing–Review & editing.

## Acknowledgments

We thank Hannah Buras for assistance with data collection.

